# Understanding Functional Roles of Native Pentose-Specific Transporters for Activating Dormant Pentose Metabolism in *Yarrowia lipolytica*

**DOI:** 10.1101/195834

**Authors:** Seunghyun Ryu, Cong T. Trinh

## Abstract

Pentoses including xylose and arabinose are the second-most prevalent sugars of lignocellulosic biomass that can be harnessed for biological conversion. Although *Yarrowia lipolytica* has emerged as a promising industrial microorganism for production of high-value chemicals and biofuels, its native pentose metabolism is poorly understood. Our previous study demonstrated that *Y. lipolytica* (ATCC MYA-2613) has endogenous enzymes for D-xylose assimilation, but inefficient xylitol dehydrogenase causes *Y. lipolytica* to assimilate xylose poorly. In this study, we investigated the functional roles of native sugar-specific transporters for activating the dormant pentose metabolism in *Y. lipolytica.* By screening a comprehensive set of 16 putative pentose-specific transporters, we identified two candidates, YALI0C04730p and YALI0B00396p, that enhanced xylose assimilation. The engineered mutants YlSR207 and YlSR223, overexpressing YALI0C04730p and YALI0B00396p, respectively, improved xylose assimilation approximately 23% and 50% in comparison to YlSR102, a parent engineered strain overexpressing solely the native xylitol dehydrogenase gene. Further, we activated and elucidated a widely unknown, native L-arabinose-assimilating pathway in *Y. lipolytica* through transcriptomic and metabolic analyses. We discovered that *Y. lipolytica* can co-consume xylose and arabinose, where arabinose utilization shares transporters and metabolic enzymes of some intermediate steps of the xylose-assimilating pathway. Arabinose assimilation was synergistically enhanced in the presence of xylose while xylose assimilation was competitively inhibited by arabinose. L-arabitol dehydrogenase is the rate-limiting step responsible for poor arabinose utilization in *Y. lipolytica*. Overall, this study sheds light on the cryptic pentose metabolism of *Y. lipolytica* and further helps guide strain engineering of *Y. lipolytica* for enhanced assimilation of pentose sugars.

**IMPORTANCE:** The oleaginous yeast *Yarrowia lipolytica* is a promising industrial platform microorganism for production of high-value chemicals and fuels. For decades since its isolation, *Y. lipolytica* has often been known to be incapable of assimilating pentose sugars, xylose and arabinose, that are dominantly present in lignocellulosic biomass. Through bioinformatic, transcriptomic and enzymatic studies, we have uncovered the dormant pentose metabolism of *Y. lipolytica*. Remarkably, unlike most yeast strains that share the same transporters for importing hexose and pentose sugars, we discovered that *Y. lipolytica* possess the native pentose-specific transporters. By overexpressing these transporters together with the rate-limiting D-xylitol and L-arabitol dehydrogenases, we activated the dormant pentose metabolism of *Y. lipolytica*. Overall, this study provides a fundamental understanding of the dormant pentose metabolism of *Y. lipolytica* and guides future metabolic engineering of *Y. lipolytica* for enhanced conversion of pentose sugars to high-value chemicals and fuels.

## INTRODUCTION

Lignocellulosic biomass derived from agricultural residues, municipal solid wastes, and woody residues can be biologically upgraded to produce high-value chemicals and fuels (1). Biomass feedstocks typically contain high contents of cellulosic and hemicellulosic components up to 80 – 90% in some biomasses such as corn cobs and nut shells (2). While cellulose is a homopolymer of glucose linked by beta-1,4-glycosidic bond with cellobiose as a structural unit, hemicellulose is a heteropolymer of pentose (C5) sugars (e.g. D-xylose, L-arabinose), hexose (C6) sugars (e.g. D-glucose, D-mannose, D-galactose), and sugar acids (3). After pretreatment and saccharification, these C5 and C6 complex sugars are released as monomers, which are subsequently fermented by microorganisms to produce target products (4). For industrial biocatalysis, microbial catalysts capable of co-utilizing all complex sugars efficiently without carbon catabolite repression are desirable.

*Yarrowia lipolytica,* a generally-regarded-as-safe (GRAS) oleaginous yeast, has emerged as a biomanufacturing platform for production of high-value chemicals and biofuels (5, 6). It has an efficient and robust native metabolism to produce high levels of organic acids and neutral lipids. Combining metabolic engineering and fermentation optimization, engineered *Y. lipolytica* strains have been generated for high production of citric acid (titer: 101.0 g/L, rate: 0.42 g/L/h; yield: 0.89 g/g glucose) (5), alpha-ketoglutaric acid (186 g/L, 1.75 g/L/h, 0.36 g/g glycerol) (6), succinic acid (160.2 g/L, 0.4 g/L/h, 0.4 g/g crude glycerol) (7), and neutral lipids (84.5 g/L, 0.73 g/L/h, 0.20 g/g glucose) (8). In addition, *Y. lipolytica* exhibits exceptional robustness to tolerate in inhibitory environments associated with biomass pretreatments. For instance, ionic liquids, 1-ethyl-3-methyl imidazolium acetate [EMIM][OAc], are promising solvents for pretreatment of recalcitrant lignocellulosic biomass (9), but are very inhibitory to microbial health at a level as low as 5% (v/v) (10). Remarkably, native *Y. lipolytica* can thrive and produce high-yield (>90%) alpha-ketoglutaric acid from cellulose in media containing up to 10% (v/v) [EMIM][OAc] (11). By screening genetic diversity from a large collection of 45 *Y. lipolytica* species isolated from different habitats, certain *Y. lipolytica* can thrive in 90% undetoxified, dilute acid-pretreated switchgrass hydrolysate and yield a relatively high lipid production (12).

One current limitation is that native *Y. lipolytica* is very inefficient at utilizing pentose sugars available from biomass hydrolysates. Figure 1 shows the pentose-assimilating pathways in yeasts. Xylose is degraded via the oxidoreductase pathway. Upon being transported intracellularly, xylose reductase (XYL1) converts xylose into D-xylitol, which is further transformed into D-xylulose by xylitol dehydrogenase (XYL2). Next, xylulokinase (XYL3) converts D-xylulose into D-xylulose-5-phosphate, which enters the pentose phosphate pathway (PPP) and is further assimilated for cell growth. On the other hand, L-arabinose is first reduced into L-arabitol by NAD(P)H-dependent arabinose reductase (ARD), which is then converted into L-xylulose by NAD(P)^+^-dependent arabitol dehydrogenase (ADH). L-xylulose is then converted to D-xylitol by NAD(P)H-dependent xylulose reductase (XLR) (13), which is further assimilated to D-xylulose-5-phosphate, a precursor for PPP. While *Y. lipolytica* is well known for assimilating hexose sugars (e.g, glucose), its pentose metabolism is dormant and poorly understood.

**Figure 1.**
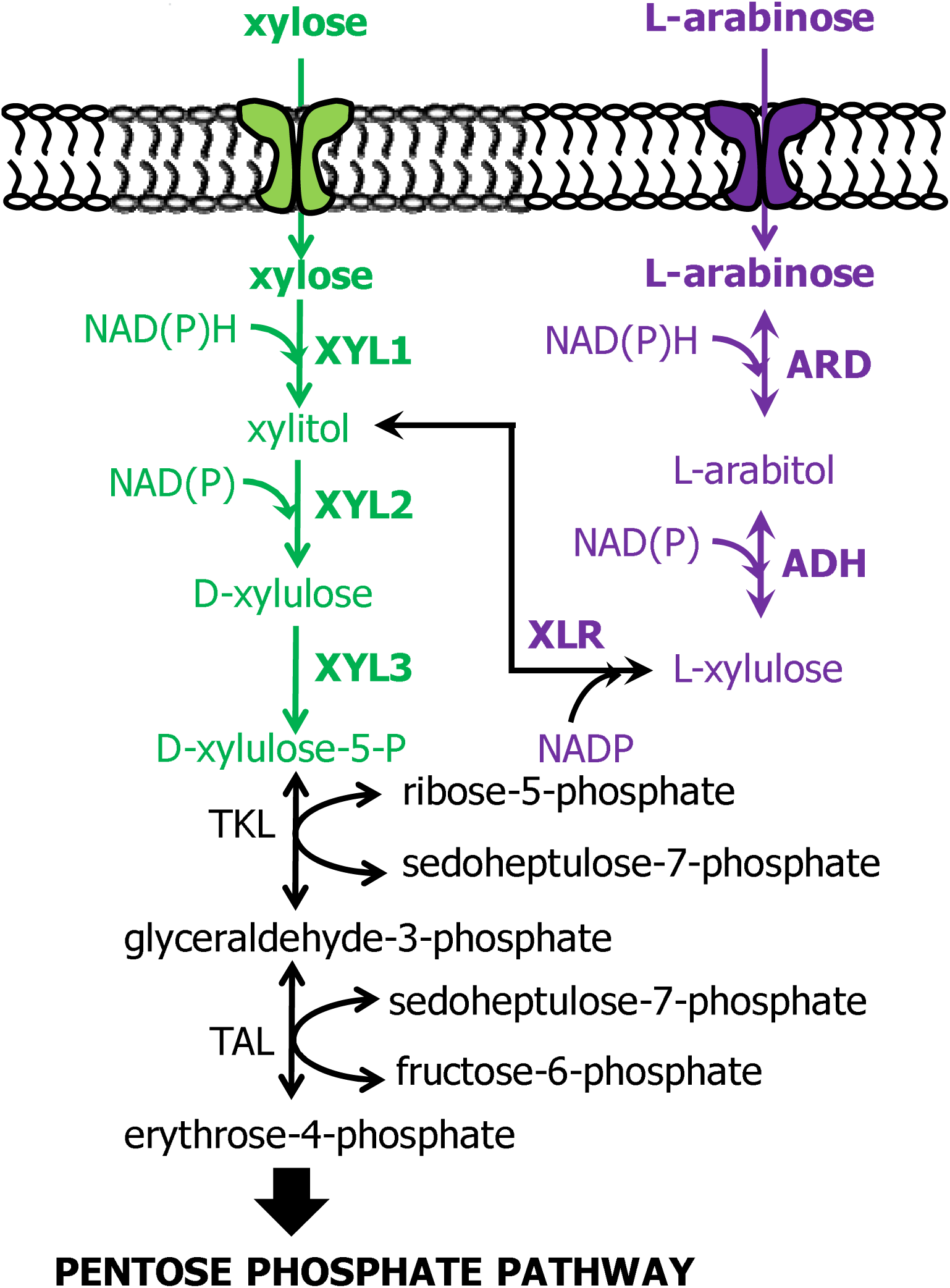
D**-**xylose (green) and L-arabinose (purple) assimilation pathways in yeast. XYL1: xylose reductase, XYL2: xylitol dehydrogenase, XYL3: xylulokinase, ARD: arabinose reductase, ADH: arabitol dehydrogenase, XLR: xylulose reductase.

We have recently activated and elucidated the native metabolism of *Y. lipolytica* for xylose assimilation. By performing bioinformatic, enzymatic, and transcriptomic analyses, we identified 16 putative pentose-specific transporters (14) and demonstrated that XYL2 is the rate-limiting step that causes poor xylose assimilation in native *Y. lipolytica*. Several metabolic engineering approaches have been applied to enhance xylose assimilation of *Y. lipolytica*. The first strategy is to express the native metabolic enzymes, e.g. XYL2 (14) or a combination of XYL2 and XYL3 (15), to improve xylose assimilation. The second approach is to overexpress heterologous enzymes of the xylose-assimilating pathway. By overexpressing *xyl1* and *xyl2* genes from *Pichia stipitis* together with implementing a starvation adaptation strategy, the engineered *Y. lipolytica* produced 15 g/L of lipid at a rate of 0.19 g/L/hr from xylose (16). Likewise, Ledesma-Amaro *et al.* generated a recombinant *Y. lipolytica* by overexpressing *xyl1* and *xyl2* genes from *P. stipitis* and *xyl3* from *Y. lipolytica*, which was able to consume 30 g/L xylose within 3 days (17) (Supplementary Table 4). It remains unknown whether *Y. lipolytica* has a native L-arabinose metabolism and is capable of assimilating this substrate as a sole carbon source for growth.

In this study, we analyzed functional roles of native pentose-specific transporters for activating a dormant pentose metabolism in *Y. lipolytica*. We screened a set of 16 putative pentose-specific transporters to identify the best candidates to enhance xylose assimilation and demonstrated that these transporters are specific to not only xylose but also arabinose. We further activated and elucidated the native arabinose-assimilating pathway in *Y. lipolytica* and shed light on functional roles of transporters and metabolic enzymes for co-utilization of xylose and arabinose. With targeted enzymatic and transcriptomic analyses, we further identified the arabitol dehydrogenase as the rate-limiting step that causes poor arabinose assimilation in *Y. lipolytica*.

## MATERIALS and METHODS

### Plasmids and strains

Table 1 shows the list of plasmids and strains used in this study and Supplementary Table 1 contains the list of primers used to construct these plasmids and strains. The plasmid, pSR008, was constructed by replacing the leucine selection marker gene (LEU2) of the plasmid pSR001 (11) with the uracil selection marker gene (URA3). The URA3 gene was amplified from the genomic DNA (gDNA) of *Y. lipolytica* NRRL YB-423 using the primers URA3_Fwd/URA3_Rev and then Gibson assembled with the backbone amplified from pSR001 using the primers pSR008_Fwd/pSR008_Rev (18).

**Table 1.**
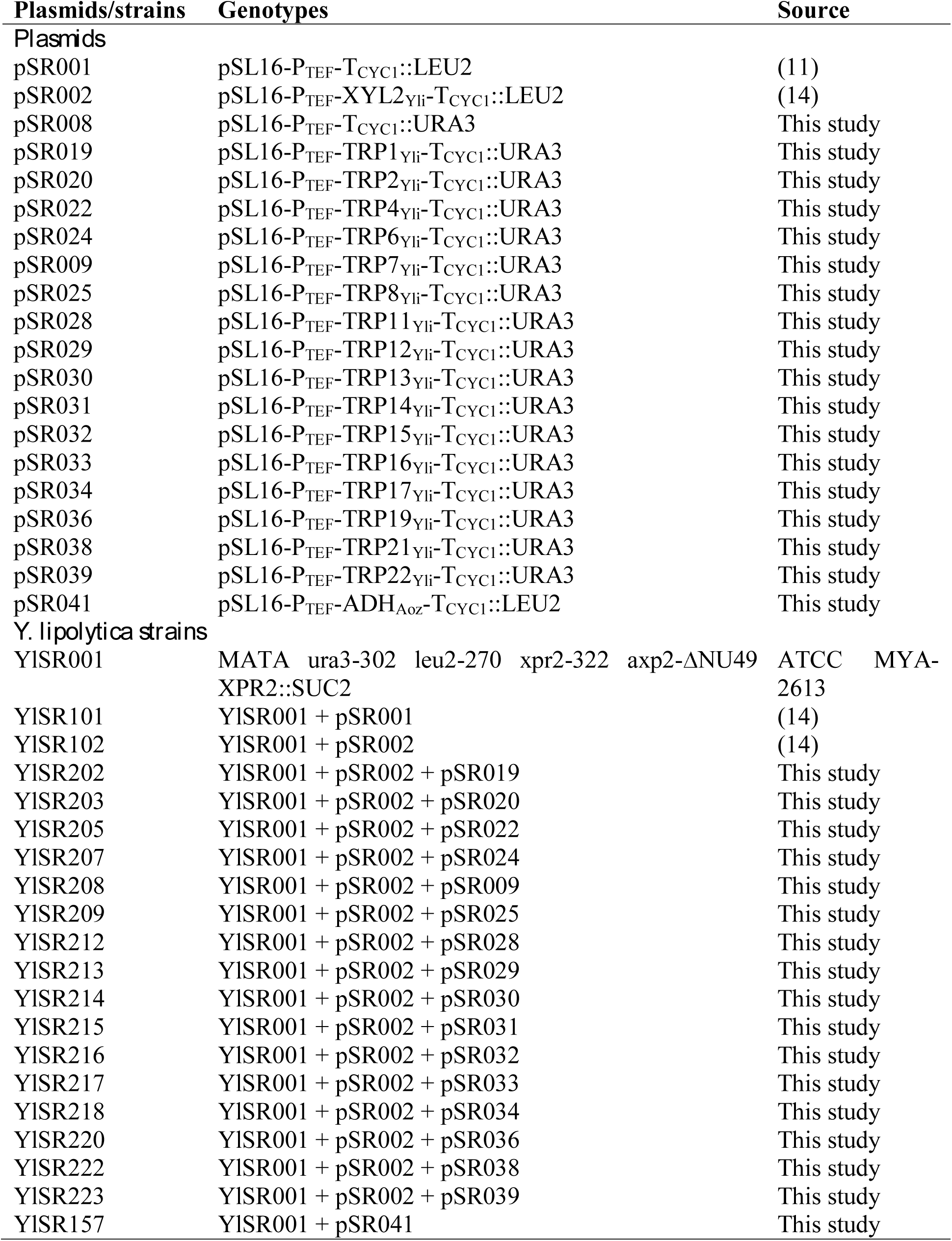
List of plasmids and strains used in this study.

| Plasmids/strains | Genotypes | Source |
| --- | --- | --- |
| *Plasmids* |  |  |
| pSR001 | pSL16-P<sub>TEF</sub>-T<sub>CYC1</sub>::LEU2 | (11) |
| pSR002 | pSL16-P<sub>TEF</sub>-XYL2<sub>Yli</sub>-T<sub>CYC1</sub>::LEU2 | (14) |
| pSR008 | pSL16-P<sub>TEF</sub>-T<sub>CYC1</sub>::URA3 | This study |
| pSR019 | pSL16-P<sub>TEF</sub>-TRP1<sub>Yli</sub>-T<sub>CYC1</sub>::URA3 | This study |
| pSR020 | pSL16-P<sub>TEF</sub>-TRP2<sub>Yli</sub>-T<sub>CYC1</sub>::URA3 | This study |
| pSR022 | pSL16-P<sub>TEF</sub>-TRP4<sub>Yli</sub>-T<sub>CYC1</sub>::URA3 | This study |
| pSR024 | pSL16-P<sub>TEF</sub>-TRP6<sub>Yli</sub>-T<sub>CYC1</sub>::URA3 | This study |
| pSR009 | pSL16-P<sub>TEF</sub>-TRP7<sub>Yli</sub>-T<sub>CYC1</sub>::URA3 | This study |
| pSR025 | pSL16-P<sub>TEF</sub>-TRP8<sub>Yli</sub>-T<sub>CYC1</sub>::URA3 | This study |
| pSR028 | pSL16-P<sub>TEF</sub>-TRP11<sub>Yli</sub>-T<sub>CYC1</sub>::URA3 | This study |
| pSR029 | pSL16-P<sub>TEF</sub>-TRP12<sub>Yli</sub>-T<sub>CYC1</sub>::URA3 | This study |
| pSR030 | pSL16-P<sub>TEF</sub>-TRP13<sub>Yli</sub>-T<sub>CYC1</sub>::URA3 | This study |
| pSR031 | pSL16-P<sub>TEF</sub>-TRP14<sub>Yli</sub>-T<sub>CYC1</sub>::URA3 | This study |
| pSR032 | pSL16-P<sub>TEF</sub>-TRP15<sub>Yli</sub>-T<sub>CYC1</sub>::URA3 | This study |
| pSR033 | pSL16-P<sub>TEF</sub>-TRP16<sub>Yli</sub>-T<sub>CYC1</sub>::URA3 | This study |
| pSR034 | pSL16-P<sub>TEF</sub>-TRP17<sub>Yli</sub>-T<sub>CYC1</sub>::URA3 | This study |
| pSR036 | pSL16-P<sub>TEF</sub>-TRP19<sub>Yli</sub>-T<sub>CYC1</sub>::URA3 | This study |
| pSR038 | pSL16-P<sub>TEF</sub>-TRP21<sub>Yli</sub>-T<sub>CYC1</sub>::URA3 | This study |
| pSR039 | pSL16-P<sub>TEF</sub>-TRP22<sub>Yli</sub>-T<sub>CYC1</sub>::URA3 | This study |
| pSR041 | pSL16-P<sub>TEF</sub>-ADH<sub>Aoz</sub>-T<sub>CYC1</sub>::LEU2 | This study |
| *Y. lipolytica strains* |  |  |
| YISR001 | MATA ura3-302 leu2-270 xpr2-322 axp2-ΔNU49 XPR2::SUC2 | ATCC MYA-2613 |
| YISR101 | YISR001 + pSR001 | (14) |
| YISR102 | YISR001 + pSR002 | (14) |
| YISR202 | YISR001 + pSR002 + pSR019 | This study |
| YISR203 | YISR001 + pSR002 + pSR020 | This study |
| YISR205 | YISR001 + pSR002 + pSR022 | This study |
| YISR207 | YISR001 + pSR002 + pSR024 | This study |
| YISR208 | YISR001 + pSR002 + pSR009 | This study |
| YISR209 | YISR001 + pSR002 + pSR025 | This study |
| YISR212 | YISR001 + pSR002 + pSR028 | This study |
| YISR213 | YISR001 + pSR002 + pSR029 | This study |
| YISR214 | YISR001 + pSR002 + pSR030 | This study |
| YISR215 | YISR001 + pSR002 + pSR031 | This study |
| YISR216 | YISR001 + pSR002 + pSR032 | This study |
| YISR217 | YISR001 + pSR002 + pSR033 | This study |
| YISR218 | YISR001 + pSR002 + pSR034 | This study |
| YISR220 | YISR001 + pSR002 + pSR036 | This study |
| YISR222 | YISR001 + pSR002 + pSR038 | This study |
| YISR223 | YISR001 + pSR002 + pSR039 | This study |
| YISR157 | YISR001 + pSR041 | This study |

A set of 16 plasmids were built containing 16 putative xylose transporter genes identified in previous paper (14). Specifically, these transporter genes were amplified from gDNA of YlSR001 using the primer sets described in Supplementary Table 1 and Gibson assembled with the pSR008 backbone amplified using the primers pSR001_Fwd/pSR001_Rev. Gene locus information for the 16 xylose-specific transporters of native *Y. lipolytca* are available in Supplementary Table 2. To construct pSR041, we used the heterologous L-arabitol dehydrogenase gene of *Aspergillus oryzae* (GenBank: BAC81768.1) that was codon optimized and gBlock synthesized by IDT (Integrated DNA Technologies, Inc. Coralville, IA), and then Gibson assembled with the pSR001 backbone (14) amplified using the primers pSR001_Fwd/pSR001_Rev. The codon optimized sequence of ADH_Aoz_ is presented in Supplementary Text 1.

*Escherichia coli* TOP10 was used for molecular cloning. All *Y. lipolytica* mutants (Table 1) were constructed by transforming YlSR001, a thiamine, leucine, and uracil auxotroph with the target plasmids via electroporation (19). All constructed *Y. lipolytica* mutants were screened and maintained in SC-Leu-Ura agar plate containing 10 g/L xylose as a carbon source. Mutants were verified by performing yeast-colony PCR (polymerase chain reaction) using the TEF promoter binding forward primer (P_TEF__seq Fwd) and gene binding reverse primer (TRPX_Yli__Rev) (Supplementary Table 1).

### Media and cell culturing

#### Media preparation

The lysogeny broth (LB) medium containing 5 g/L yeast extract, 10 g/L tryptone, and 5 g/L NaCl was used for molecular cloning in *E. coli.* For selection, 100 µg/mL ampicillin was added in LB medium. For *Y. lipolytica* characterization experiments, the synthetic SC-Leu-Ura media (pH 5.5) were used containing yeast nitrogen base (cat no. Y0626, Sigma-Aldrich, MO, USA), synthetic dropout amino acids mixture without leucine, uracil, and tryptophan (cat no. Y1771, Sigma-Aldrich, MO, USA; or cat no. AC172110250, Fisher Scientific, MA, USA), L-tryptophan (cat no. AC140590010, Acros Organics, NJ, USA), and 30 µg/mL chloramphenicol. Single or mixture of sugars, including glucose, xylose, and/or arabinose, were also added to the yeast synthetic media at an initial concentration of 10 g/L.

#### Isolation of efficient xylose-utilizing mutants

To isolate the efficient xylose-utilizing mutants, each of the 16 *Y. lipolytica* mutants, YlSR202, YlSR203, YlSR205, YlSR207-209, YlSR212-218, YlSR220, YlSR222, and YlSR223, carrying the native xyl2 and putative xylose transporter genes, were first grown in a 15 mL culture tube containing 1 mL SC-Leu-Ura with 10 g/L glucose as a carbon source. When cell growth reached a mid-exponential phase (optical density(OD) at 600nm ∼ 3), cells were spun down and the cell pellet was washed with 5 mL sterile water three times to remove any residual glucose. Next, the pellet was resuspended in the SC-Leu-Ura medium without sugars and diluted 10x, 100x, and 1000x with the same medium. The diluted cultures of each mutant were then spotted on the SC-Leu-Ura plate containing 10 g/L xylose. Mutants exhibiting fast growth on xylose were selected for subsequent characterization studies.

#### Cell culturing

Growth characterization of *Y. lipolytica* in the yeast synthetic media, containing single and a mixture of xylose and/or arabinose, was conducted in a 500 mL baffled flask with a 50 mL working volume at 28^o^C and 300 rpm. The initial OD was adjusted to 0.2. Time profiles of cell growth and metabolites were measured during cell culturing and used to determine kinetic parameters, such as specific cell growth rate (μ, 1/hr) and specific sugar uptake rate (r_S_, mmol/g DCW/hr) (20). The gene expression and enzyme activity were also measured for cell cultures collected during the mid-exponential growth phase. All experiments were performed with at least 6 biological replicates.

### Bioinformatics

BlastP (21) was applied to identify the putative L-arabinose reductase (ARD), L-arabitol dehydrogenase (ADH), and L-xylulose reductase (XLR) enzymes of the arabinose-assimilating pathway of *Y. lipolytica*. We used ARD_Ani_ (from *Aspergillus niger*, uniprot: A2QBD7 (22)), ADH_Ncr_ (from *Neurospora crassa*, uniport: Q7SI09 (23)), NADPH-dependent XLR_Ani_ (uniport: G3YG17 (24)), NADPH-dependent XLR_Tre_ (from *Trichoderma reesei*, uniport: G0RH19) (22)), and NADH-dependent XLR_Amo_ (from *Ambrosiozyma monospora*, uniport: Q70FD1_9ASCO (25)) as BlastP templates.

### Analytical methods

#### High-performance liquid chromatography (HPLC)

Extracellular metabolites (e.g, sugars and sugar alcohols) were analyzed using a Shimadzu HPLC system, equipped with a refractive index detector (RID) (Shimadzu Scientific Instruments, Inc., MD, USA). Prior to HPLC run, culture medium was filtered through a 0.2 μm pore size membrane filter. To analyze glucose, we used a Biorad Aminex 87H column (cat no. 1250140, Bio-Rad Laboratories, CA, USA) set at 48°C and 10 mN H_2_SO_4_ as a mobile phase operating at 0.6 mL/min (14). To analyze xylose, xylitol, arabinose, and arabitol, we used a Biorad Aminex 87C column (cat no. 1250095, Bio-Rad Laboratories, CA, USA) set at 85°C and water as a mobile phase running at 0.6 mL/min.

#### Transcriptomics by real time-PCR (rt-PCR)

Gene expression levels were quantified using rt-PCR. First, mid-exponential growth phase cells (OD in a range of 2.0–2.5) were harvested. Total RNA was purified using a Qiagen RNeasy mini kit (Cat no. 74104, Qiagen Inc, CA, USA.), and cDNA was synthesized using a QuantiTect Reverse Transcription kit (Cat no. 205311, Qiagen Inc, CA, USA.). The rt-PCR experiment was performed using a QuantiTect SYBR Green PCR kit (Cat no. 204143, Qiagen Inc, CA, USA) and a StepOnePlus™ Real-Time PCR System (Applied Biosystems, CA, USA). A relative mRNA expression level of a target gene at a given condition was normalized to that of a house-keeping actin gene (YALI0D08272g) as previously described (14). Primers used for rt-PCR are listed in Supplementary Table 1. To compare relative gene expression of a target gene under two different conditions, we calculated the log2 ratio of the expression levels for that target gene between condition 1 (e.g., growth on arabinose) and condition 2 (e.g., growth on xylose as a reference condition) (26). A relative mRNA expression level for each gene under a given growth condition was reported as an average ± 1 standard deviation from a data set of at least three biological replicates. The students *t*-test was performed to evaluate statistical significance.

### *In vitro* enzyme activity assays

To prepare cell lysates, cell cultures were collected during the mid-exponential growth phase (OD between 2.0 and 2.5) and suspended in a Y-PER yeast protein extraction reagent (cat no. 78990, Thermo Scientific, IL, USA) including 2× EDTA-free pierce protease inhibitors (cat no. 88266 Thermo Scientific, IL, USA). Cells were then lyzed by incubation at 28°C, 300 rpm for 60 min. The soluble fraction was separated by centrifugation at 17,000xg for 10 min and used for enzyme activity assays.

Each *in vitro* enzyme activity assay for arabinose reductase (ARD), xylitol dehydrogenase (XYL2), arabitol dehydrogenase (ADH), and xylulose reductase (XLR) was conducted by adding 1.0 µg of whole-cell lysate in a reaction mixture containing 25 mM sodium phosphate buffer (pH 6.0), appropriate cofactor (e.g. NAD(P)H or NAD(P)^+^), and substrate (e.g. L-arabinose, D-xylitol, L-arabitol, or L-xylulose) in 384-well plates with a 70 µl working volume at 28°C. For the ARD assay, 1 mM NAD(P)H and 300 mM L-arabinose were used, and the reduction rate of NAD(P)H was monitored at 340 nm. For the XYL2 and ADH assays, 1 mM NAD(P)^+^ and 300 mM substrate (i.e. xylitol for the XYL2 and arabitol for ADH) were used. For the XLR activity assay, 1 mM NAD(P)H and 30 mM L-xylulose were used. One unit of each enzyme activity was defined as one µmole of NAD(P)H generated or reduced per mg protein per min.

Enzyme kinetics was measured using a BioTek Synergy HT microplate reader with an associated Gen5 software (BioTek Instruments, Inc., VT, USA). Protein concentration was quantified by the Bradford method (27). All enzyme assay experiments were performed with at least three biological replicates.

## RESULTS AND DISCUSSION

### Screening putative pentose-specific transporters for enhanced xylose consumption

We constructed a set of 16 *Y. lipolytica* strains, YlSR202, YlSR203, YlSR205, YlSR207-YlSR209, YlSR212-YlSR218, YlSR220, YlSR222, and YlSR223, each of which overexpresses a putative pentose-specific transporter and a native xylitol dehydrogenase (Table 1). To identify the best xylose-assimilating candidates, we screened these strains for fast growth on solid agar plates containing xylose as a carbon source. The result shows that YlSR207 and YlSR223, carrying TRP6_Yli_ (YALI0C04730g) and TRP22_Yli_ (YALI0B00396g), respectively, exhibited enhanced growth on xylose within 72 hr incubation, while other *Y. lipolytica* strains exhibited either poor or no growth (Figure 2). Poor growth may be attributed to low affinity of the transporters toward xylose, improper folding of overexpressed transporters, and/or incorrect membrane localization.

**Figure 2.**
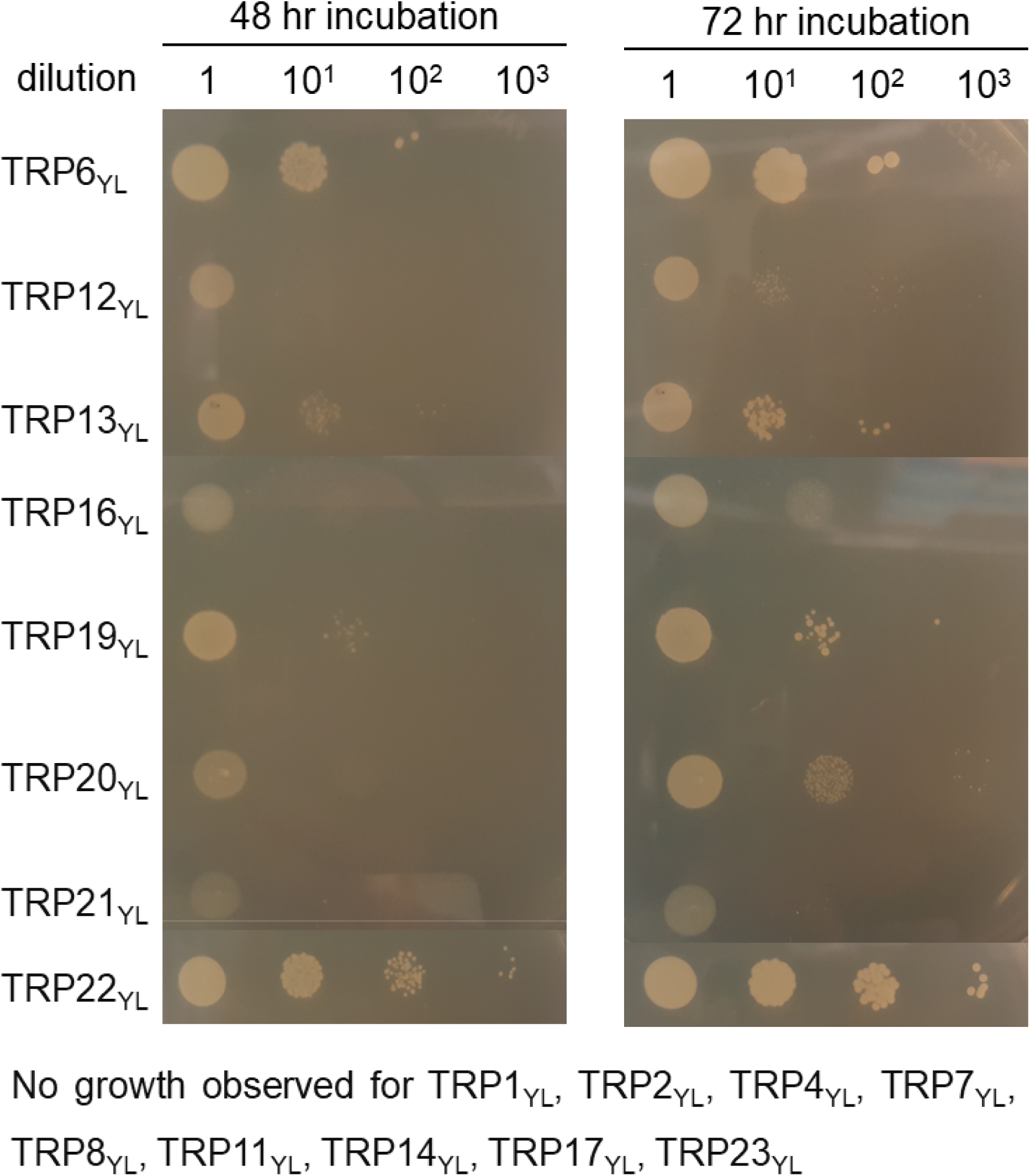
Growth screening of *Y. lipolytica* strains expressing putative xylose-specific transporters on SC-Leu-Ura agar plate containing 10 g/L xylose as a single sugar.

Overall, we identified the two *Y. lipolytica* strains, YlSR207 carrying TRP6_Yli_ and YlSR223 carrying TRP22_Yli_, exhibited the most effective growth on xylose. This result provides a basis for detailed characterization of functional roles of these transporters for activating the native pentose metabolism in *Y. lipolytica* in subsequent studies.

### Elucidating TRP6_Yli_ and TRP22_Yli_ are xylose-specific transporters in *Y. lipolytica*

To demonstrate TRP6_Yli_ and TRP22_Yli_ are xylose-specific transporters *in vivo*, we characterized YlSR207 and YlSR223 in a defined, xylose-containing SC-Leu-Ura liquid medium. As a reference, we chose YlSR202 that overexpressed TRP1_Yli_ and did not improve xylose assimilation (Figure 2). The result shows that YlSR202, YlSR207, and YlSR223 grew on xylose as a single carbon source without significant accumulation of xylitol (Figure 3A and3B) because they overexpressed XYL2_Yli_ that has been previously shown as the rate-limiting step of xylose-assimilating pathway in *Y. lipolytica* (14). Specifically, the reference strain YlSR202 grew most slowly with a specific growth rate of 0.014 ± 0.001 (1/hr) and took 192 hr to completely consume 10 g/L xylose with a specific xylose uptake rate of 0.154 ± 0.014 (mmol/gCDW/hr) (Table 2). YlSR202 exhibited the same growth phenotype as YlSR102 that overexpressed only XYL2_Yli_ without any transporter as previously characterized (14). In contrast, YlSR207 (0.019 ± 0.004 1/hr) and YlSR223 (0.023 ± 0.001 1/hr) grew 40% and 62% faster than YlSR202, respectively, under the same growth condition (*p*-value < 0.05, n ≥ 6). Both YlSR207 and YlSR223 were also capable of completely consuming 10 g/L xylose within 108 hr with a 23% and 50% increase in the specific xylose uptake rates, respectively (*p*-value < 0.05, n ≥ 6).

**Figure 3.**
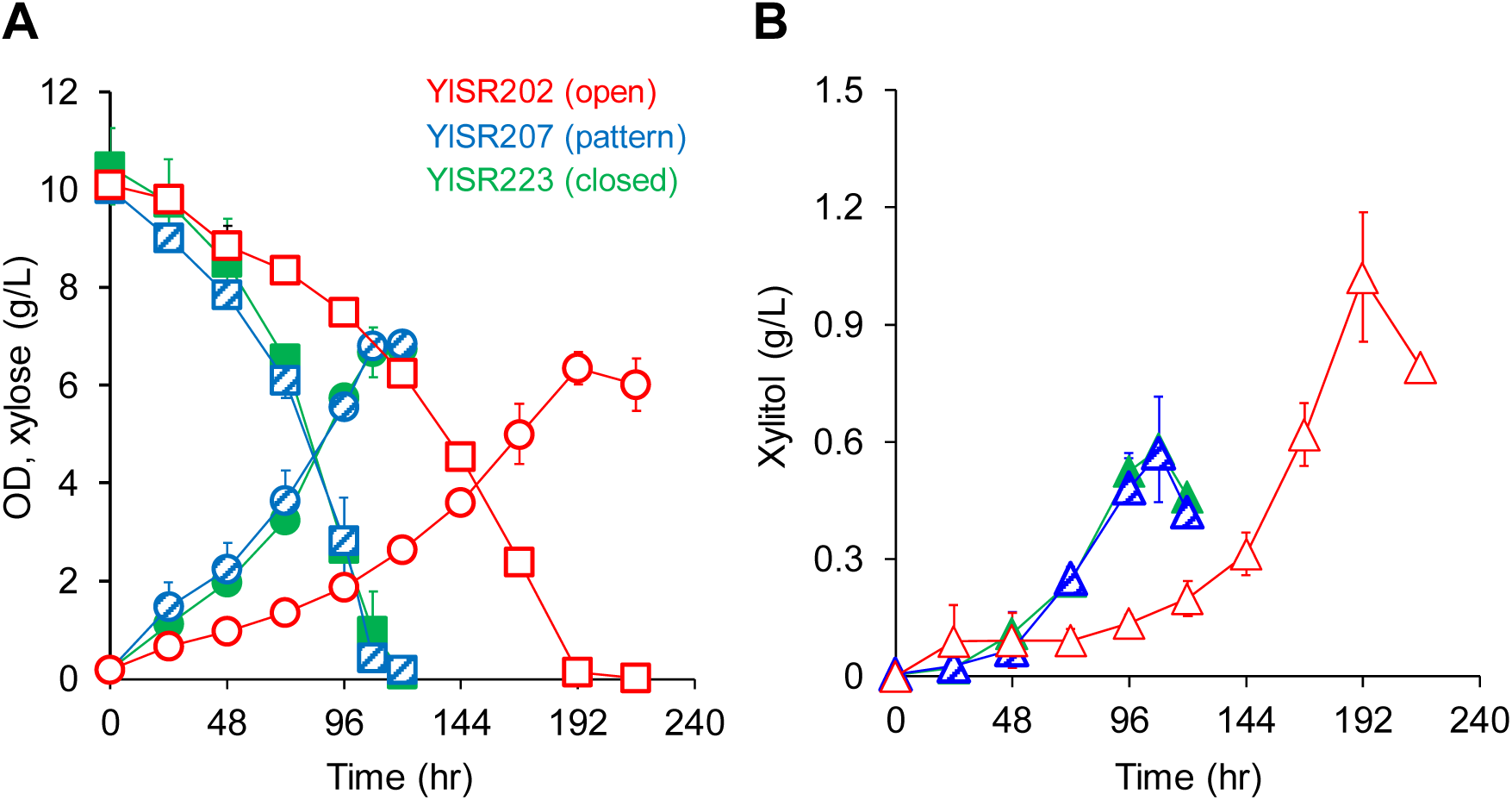
Profiles of cell growth and metabolites of *Y. lipolytica* strains, YlSR202, YlSR207, and YlSR223, growing on single xylose. Symbols: OD (circle), xylose (square), and xylitol (triangle). Each data point represents an average value ± standard deviation from the results of 6 biological replicates.

**Table 2.** Specific growth rates (μ, 1/hr) and specific sugar uptake rates (r_S_, mmol/gCDW/hr) of *Y. lipolytica* strains, YlSR102, YlSR207, and YlSR223, growing on single xylose (xyl) and mixed pentose sugars (xyl + ara).

| Strains | $\mu$ | | $r_{xyl}$ | | $r_{ara}$ |
| --- | --- | --- | --- | --- | --- |
|  | xyl | xyl + ara | xyl | xyl + ara | xyl + ara |
| YISR102 | 0.014±0.001 | 0.007±0.001 | 0.154±0.014 | 0.095±0.007 | 0.070±0.007 |
| YISR207 | 0.019±0.004 | 0.018±0.000 | 0.190±0.039 | 0.170±0.028 | 0.043±0.014 |
| YISR223 | 0.023±0.001 | 0.019±0.001 | 0.231±0.042 | 0.163±0.011 | 0.021±0.006 |

To further validate that the improvement on cell growth and xylose consumption of YlSR207 and YlSR223 are due to overexpression of TRP6_Yli_ and TRP22_Yli_ but not XYL2_Yli_, we showed that the strains YlSR202, YLSR207, and YlSR223 exhibited the same xylitol dehydrogenase XYL2 activity regardless of cofactors (NAD^+^ or NADP^+^) used for the assay (*p*-value < 0.05, n ≥ 6) (Supplementary Figure 1). By reviving YlSR207 and YlSR223 strains from frozen glycerol stocks, we confirmed their stable growth phenotypes since their growth rates remained unaffected. Further, YlSR207 and YlSR223 exhibited the same growth rates without lag when they were first cultured in glucose media and then transferred to fresh xylose media.

Taken altogether, *Y. lipolytica* has not only native metabolic enzymes (14) but also xylose-specific transporters for xylose assimilation as demonstrated here even though its xylose metabolism stays dormant due to transcriptional repression. This native xylose metabolism can be activated without a need to express any heterologous enzyme. We demonstrated for the first time that TRP6_Yli_ and TRP22_Yli_ are native xylose-specific transporters of *Y. lipolytica*. In yeast, specificity of xylose transporter is determined by “G-G/F-X-X-X-G” structural motif and two conserved amino acids, threonine and asparagine (i.e. T213 and N370 in *S. cerevisiae* HXT7) (28, 29). TRP6_Yli_ (YALI0C04730p) contains a “G-F-L-L-F-G” structural motif and tyrosine instead of asparagine (N348Y) (14). Although no conserved structural motif is found in TRP22_Yli_ (YALI0B00396p), valine, a small hydrophobic amino acid, instead of threonine and asparagine contributes to the xylose specificity (14).

Engineering pentose-specific transporters is critical for enhanced pentose assimilation in yeasts, especially in presence of competitive glucose substrate. For instance, *S. cerevisiae* is not known to have xylose-specific transporters, therefore it is required to express the heterologous transporter *Candida intermedia* Gxf1 (30) or engineer the native glucose-specific transporter HXT7 to be xylose-specific (28, 29) for enhanced xylose assimilation. The discovery of functional roles of TRP6_Yli_ and TRP22_Yli_ will be useful for metabolic engineering of other yeasts (such as *S. cerevisiae.*) to enhance xylose assimilation.

### Understanding the native L-arabinose-assimilating pathway in *Y. lipolytica*

***Degradation of single arabinose*.** The existence and function of the native arabinose-assimilating pathway in *Y. lipolytica* is widely unknown. To investigate the native arabinose metabolism of *Y. lipolytica*, we first performed genome mining. The bioinformatic result shows that *Y. lipolytica* has the putative enzymes of the arabinose-assimilating pathway, including arabinose reductase (*ARD*), arabitol dehydrogenase (*ADH*), and xylulose reductase (*XLR*) (Figure 1). We identified 11 putative ARD genes, 5 putative XLR genes, and one putative ADH gene (see Supplementary Table 3 for their gene loci). Interestingly, the arabitol dehydrogenase ADH_Yli_ YALI0E12463g is the same gene identified as a xylitol dehydrogenase (XYL2_Yli_) gene (14).

To test whether the arabinose-assimilating pathway is active in *Y. lipolytica*, we first cultured YlSR102 (overexpressing XYL2_Yli_) in the defined liquid medium containing arabinose as a carbon source. The result shows that YlSR102 grew poorly on arabinose and only consumed insignificant amount of arabinose (Figure 4A). However, transcriptomics shows that the arabinose-assimilating pathway genes were up-regulated. Four ARD genes – *ARD6*, *ARD7*, *ARD8*, and *ARD9* out of 11 putative ARD genes-were up-regulated in cell cultures growing on single arabinose compare to cells grown in single xylose by 2.74 ± 1.40-fold, 10.77 ± 2.95-fold, 7.21 ± 1.06-fold, 5.86 ± 0.23-fold, respectively (Figure 5). Two XLR genes -*XLR1* and *XLR4* out of 5 putative XLR genes – were up-regulated by 5.13 ± 0.29-fold and 4.32 ± 0.46-fold, respectively. The ADH_Yli_ (also identified as XYL2_Yli_) gene was up-regulated by 1.81 ± 0.70-fold. Consistent with transcriptomic data, we also detected ARD, ADH, and XLR enzyme activities from YlSR102 cultures (Table 3). The ARD and XLR activities, specific to NADPH, were significantly higher for YlSR102 growing on arabinose than xylose (negative control). In contrast, the ADH activity, specific to NAD^+^, remains similar between cells grown in single arabinose and single xylose. In addition, we characterized YlSR207 and YlSR223 to investigate whether TRP6_Yli_ and TRP22_Yli_ helped enhance arabinose assimilation. However, YlSR207 and YlSR223 poorly consumed arabinose like YlSR102 (Figure 4B and 4C).

**Figure 4.**
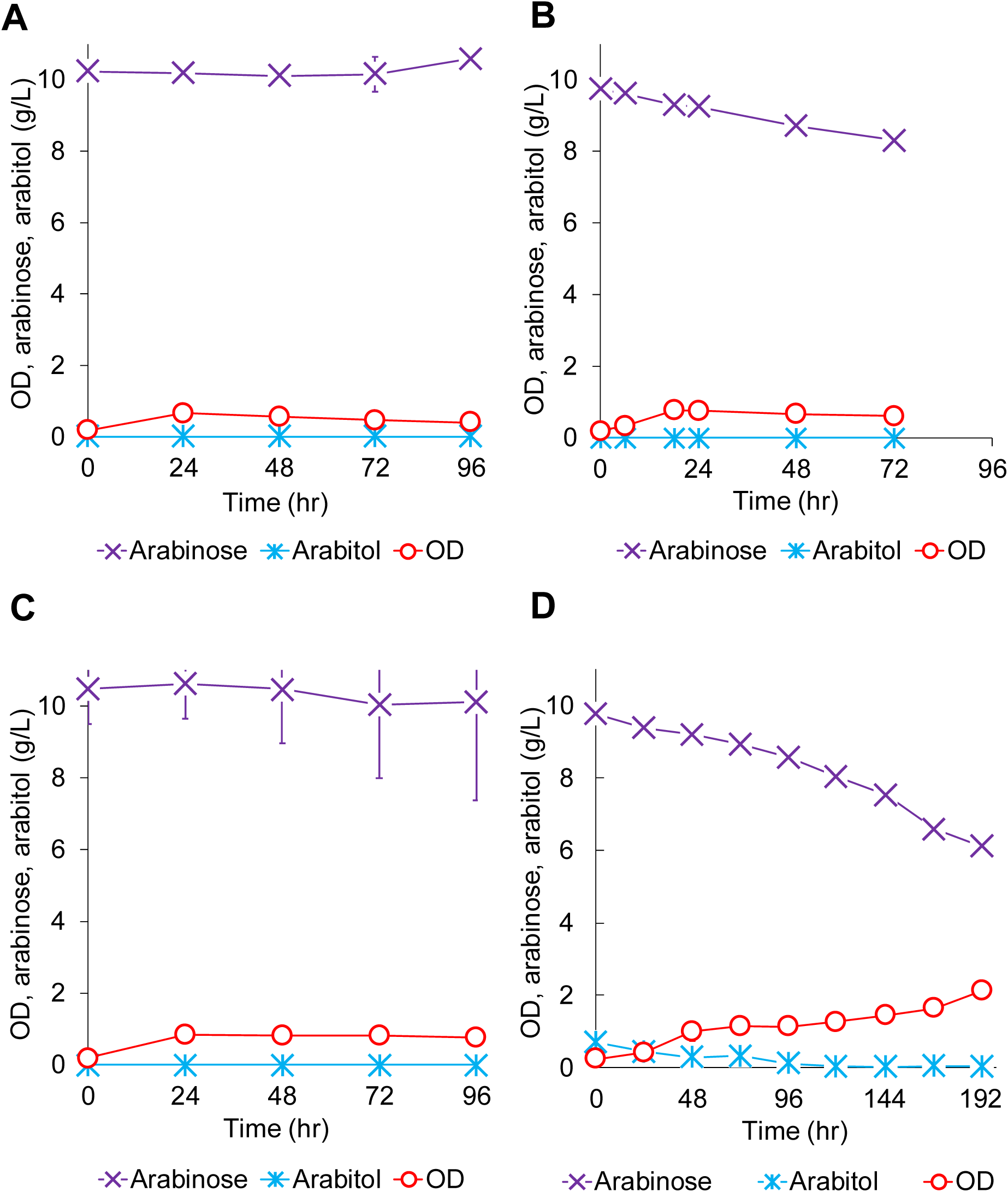
Profiles of cell growth and metabolites of *Y. lipolytica* strains, including (A) YlSR102, (B) YlSR207, (C) YlSR223, and (D) YlSR157 growing on single arabinose. Each data point represents an average value ± 1 standard deviation from the results of 6 biological replicates.

**Figure 5.**
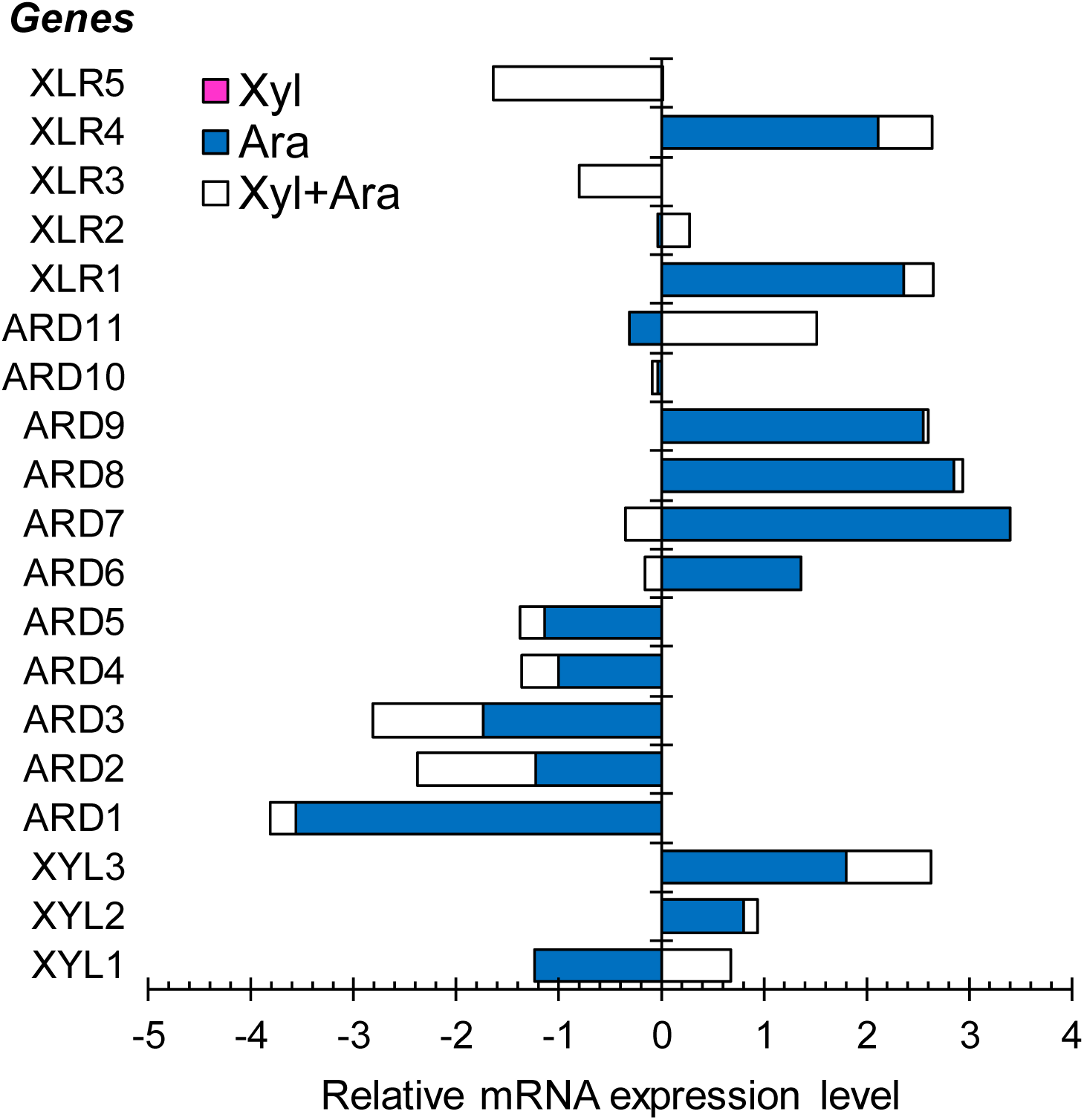
Relative mRNA expression levels (log2 scale) of putative pentose-assimilating pathway genes of *Y. lipolytica* YlSR102 growing on single xylose (xyl), single arabinose (ara), and mixed pentose sugars (xyl+ara). The reference condition for normalization is growth on single xylose. Each data point represents an average value of 3 biological replicates. Error data are not included in the figure due to crowding effect but presented in Supplementary Table 3.

**Table 3.**
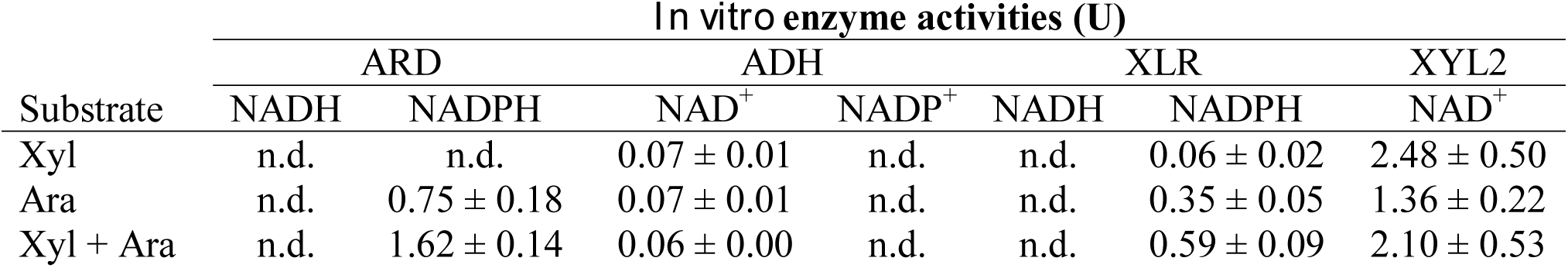
*In vitro* enzyme activities (U) of arabinose reductase (ARD), arabitol dehydrogenase (ADH), xylulose reductase (XLR), and xylitol dehydrogenase (XYL2) of YlSR102 growing on single xylose (xyl), single arabinose (ara), and mixed pentose sugars (xyl + ara). NAD(P)^+^ and NAD(P)H were used as cofactors for enzyme assays. Abbreviation – n.d.: not detected.

| Substrate | *In vitro* enzyme activities (U) |  |  |  |  |  |  |
| --- | --- | --- | --- | --- | --- | --- | --- |
|  | ARD |  | ADH |  | XLR |  | XYL2 |
|  | NADH | NADPH | NAD<sup>+</sup> | NADP<sup>+</sup> | NADH | NADPH | NAD<sup>+</sup> |
| Xyl | n.d. | n.d. | 0.07 ± 0.01 | n.d. | n.d. | 0.06 ± 0.02 | 2.48 ± 0.50 |
| Ara | n.d. | 0.75 ± 0.18 | 0.07 ± 0.01 | n.d. | n.d. | 0.35 ± 0.05 | 1.36 ± 0.22 |
| Xyl + Ara | n.d. | 1.62 ± 0.14 | 0.06 ± 0.00 | n.d. | n.d. | 0.59 ± 0.09 | 2.10 ± 0.53 |

Taken altogether, the results suggest that *Y. lipolytica* has endogenous metabolic capability to assimilate arabinose as a carbon source but levels of gene expression and/or activities of metabolic enzymes and transporters might not be sufficient. The cofactor imbalance, due to ARD and XLR specificity toward NADPH and ADH toward NAD^+^, might have also contributed to poor arabinose assimilation in *Y. lipolytica*.

#### Degradation of a mixture of xylose and arabinose by YlSR102

Since arabinose- and xylose-assimilating pathways are interconnected (Figure 1), especially when *Y. lipolytica* has only one putative YALI0E12463p enzyme that functions as xylitol dehydrogenase (XYL2_Yli_) and arabitol dehydrogenase (ADH_Yli_), we investigated whether activation of the native xylose metabolism of *Y. lipolytica* could enhance arabinose assimilation by growing it on a mixture of xylose and arabinose.

Characterization of YlSR102 shows that it was able to grow in a mixture of xylose and arabinose (Figure 6A). Up-regulation of the xylose- and arabinose-metabolizing genes further validated the active pentose metabolism of *Y. lipolytica* (Figure 5). YlSR102 consumed 1.82 ± 0.81 g/L arabinose with a specific arabinose uptake rate of 0.070 ± 0.007 (mmol/gCDW/hr) (Table 2) and accumulated 0.13 ± 0.00 g/L arabitol after 216 hr. Consistently, we also detected ARD, ADH, and XLR activities from YlSR102 cultures (Table 3). While the ARD and XLR activities were 2-fold higher for YlSR102 growing on mixed pentose sugars than single arabinose, the ADH activity remained almost the same. This result strongly suggests that *Y. lipolytica* has an active arabinose-assimilating pathway but arabitol dehydrogenase is likely a rate-limiting step.

**Figure 6.**
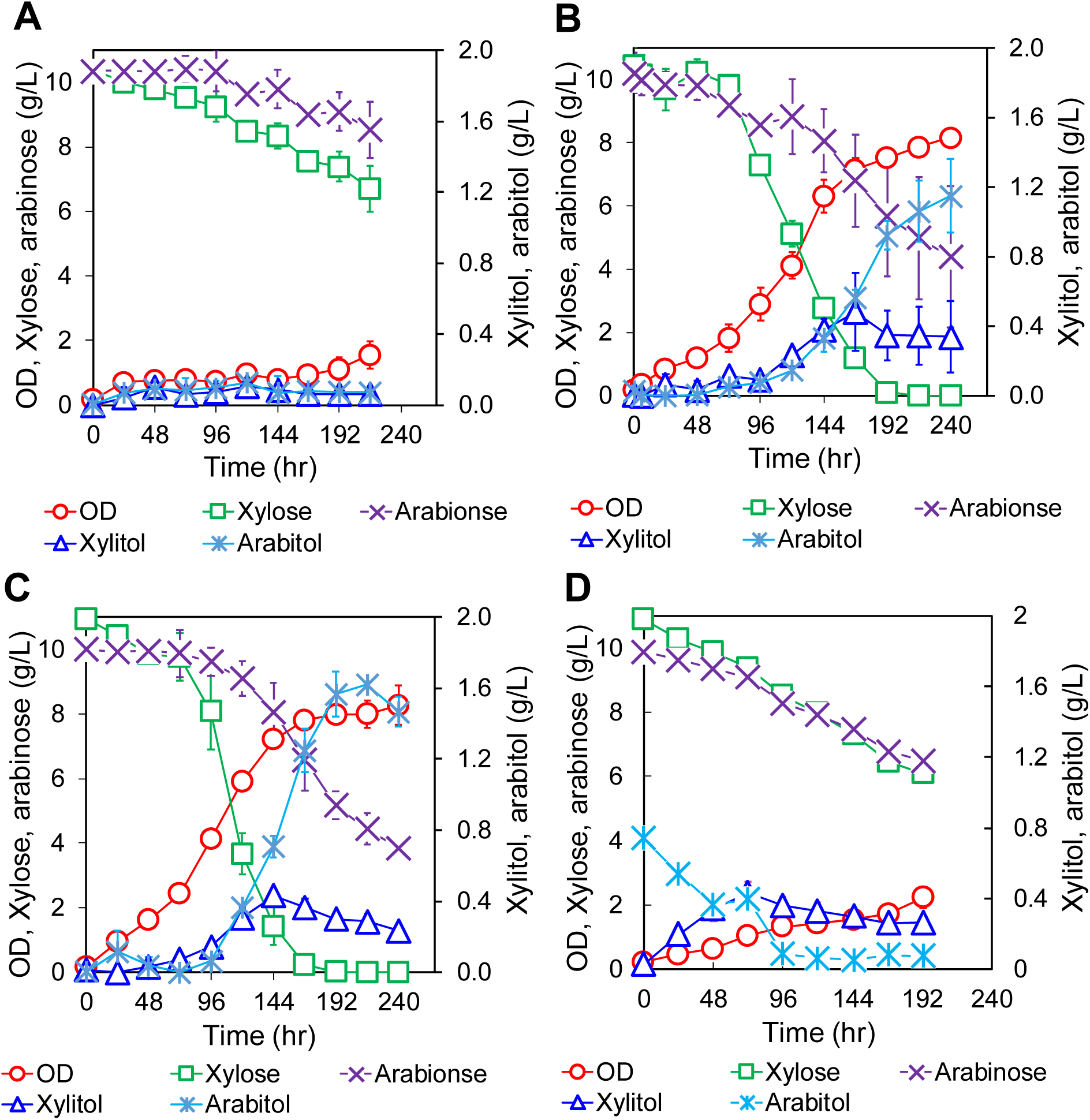
Profiles of cell growth and metabolites of *Y. lipolytica* strains, including (A) YlSR102, (B) YlSR207, (C) YlSR223, and (D) YlSR157 growing on mixed pentose sugars. Each data point represents an average value ± 1 standard deviation from the results of 6 biological replicates.

In contrast to the enhanced arabinose assimilation, YlSR102 exhibited a lower xylose consumption rate for growth on mixed pentose sugars than single xylose by 1.619 ± 0.040-fold (Table 2). To explain this phenotype, we analyzed the XYL2 and ADH activities of YlSR102 growing on single (control) and mixed sugars since XYL2_Yli_ and ADH_Yli_ were encoded by the same gene YALI0E12463g (Table 3). We observed that the XYL2 activities were relatively similar for growth on xylose and mixed pentose sugars, but lower for growth on arabinose. The ADH activities were similar under all three conditions but significantly lower than the XYL2 activities. Taken altogether, these results suggest that the competition of XYL2 might have resulted in reduced xylose consumption when YlSR102 grew on mixed pentose sugars. Poor ADH activities were also likely responsible for inefficient arabinose assimilation. In some xylose-fermenting yeasts such as *N. crassa* and *P. stipitis*, xylose reductase (XYL1) has an activity for both xylose and arabinose substrates (31, 32). Since xylose reductase of *Y. lipolytica* is 48.89% and 55.56% identical to those of *P. stiptis* (Uniprot P31867) and *N. crassa* (Uniprot Q7SD67), respectively, it is also plausible that the competition of xylose and arabinose substrates for this xylose reductase might have additionally reduced xylose assimilation efficiency.

Overall, the results show that the native arabinose metabolism of *Y. lipolytica* was active and arabinose utilization was synergistically enhanced by xylose. Xylose assimilation was however repressed by arabinose due to possible substrate competition of shared intermediate enzymes of xylose- and arabinose-assimilating pathways. L-arabitol dehydrogenase (ADH) might be a major bottleneck for efficient arabinose assimilation for growth.

#### Degradation of a mixture of xylose and arabinose by YlSR207 and YlSR223

To further investigate the functional roles of TRP6_Yli_ and TRP22_Yli_ in activating pentose metabolism, we grew YlSR207 and YlSR223 on a mixture of xylose and arabinose. The results show that YlSR207 and YlSR223 grew much faster than YlSR102 with > 2.7-fold increase in the specific growth rate and > 4.4-fold increase in biomass titer (Figure 6B-C and Table 2). We also observed that the arabinose consumptions of YlSR207 and YlSR223 was synergistically enhanced. The YlSR207 and YlSR223 consumed 5.80 ± 1.58 g/L and 6.14 ± 0.01 g/L arabinose, respectively, with 1.15 ± 0.21 g/L and 1.47 ± 0.08 g/L arabitol accumulation, respectively.

Unlike YlSR102, both YlSR207 and YlSR223 completely consumed 10 g/L xylose within 192 hr, with 1.78-fold and 1.71-fold increase in specific xylose uptake rates, respectively. The strain YlSR207 achieved very similar specific xylose uptake rates when growing on either single xylose or mixed pentose sugars (Table 2). In contrast, the xylose specific uptake rate of YlSR223 was 1.40-fold lower for growth on mixed pentose sugars than single xylose. Since arabinose assimilation is xylose-dependent, it is difficult to directly compare the efficiency of TRP6_Yli_ and TRP22_Yli_ to enhance arabinose assimilation. To examine this, we first cultured YlSR207 and YlSR223 in single xylose (phase 1) until substrate completion, followed by addition of 10 g/L arabinose (phase 2) (Supplementary Figure 2). The results show that there is no statistical significance on the specific arabinose consumption rates for YlSR207 (0.068 ± 0.020 mmol/gCDW/hr) and YlSR223 (0.042 ± 0.005 mmol/gCDW/hr), respectively.

Taken altogether, the results strongly validated the existence of the dormant pentose metabolism of *Y. lipolytica*. Overexpressing the native xylitol/arabitol dehydrogenase (XYL2_Yli_) and pentose transporters (TRP6_Yli_ and TRP22_Yli_) can activate pentose metabolism. Shared common intermediate enzymes and transporters of xylose- and arabinose-degrading pathways clearly affect the efficiency of pentose assimilation. The low activity of the endogenous arabitol dehydrogenase and cofactor imbalance likely hinder arabinose assimilation. TRP6_Yli_ and TRP22_Yli_ have *in vivo* activities toward both xylose and arabinose, and their overexpression enhance pentose assimilation. Like *Y. lipolytica*, other yeasts (except *S. cerevisiae*) including *Candida arabinofermentans* PYCC 5603T, *Pichia guilliermondii* PYCC 3012, and *Ambrosiozyma monospora* also possess arabinose transporters (33, 34).

### Alleviating the L-arabitol dehydrogenase rate-limiting step in *Y. lipolytica*

Combination of bioinformatic and enzymatic analyses and arabitol accumulation clearly suggests that L-arabitol dehydrogenase is the rate-limiting step that causes poor arabinose assimilation in *Y. lipolytica*. To truly validate this, we constructed the strain YlSR157 that overexpressed a heterologous arabitol dehydrogenase of *Aspergillus oryzae* (ADH_Aoz_) using the constitutive TEF promoter. ADH_Aoz_ was chosen because it exhibited higher activities toward L-arabitol (39.2 mU/mg protein) than D-xylitol (6.54 mU /mg protein) (35).

Unlike YlSR102, YlSR157 grew faster and consumed more sugars when growing on both single arabinose and mixed pentose sugars (Figure 4D and 6D). In single arabinose, YlSR157 reached final OD of 2.13 ± 0.02 and consumed 3.66 ± 0.13 g/L arabinose with insignificant accumulation of arabitol after 193 hr (Figure 4D). For growth on mixed pentose sugars, YlSR157 achieved a final OD of 2.23 ± 0.31 and consumed a total of 4.80 ± 0.51 g/L xylose and 3.40 ± 0.43 g/L arabinose with insignificant accumulation of xylitol (0.27 ± 0.01 g/L) and arabitol (0.08 g/L ± 0.00 g/L) after 193 hr (Figure 6D). Consistent with growth phenotypes, YlSR102 yielded higher XYL2 activity than YlSR157 by 3.1-fold (Supplementary Figure 3). While YlSR102 barely exhibited any ADH activity, YlSR157 achieved a significantly higher (>50 fold) ADH activity. These results suggest that ADH_Aoz_ exhibited activities toward both D-xylitol and L-arabitol, but has higher specificity for L-arabitol than D-xylitol. Like XYL2_Yli_, ARD_Aoz_ is specific to NAD^+^ only (data not shown). This cofactor imbalance might help explain incomplete arabinose assimilation when YlSR157 grew on either single arabinose or mixed pentose sugars.

Taken altogether, we validated that arabitol dehydrogenase is the rate-limiting step in *Y. lipolytica*. While overexpressing ADH_Aoz_ enhanced arabinose assimilation due to its higher activity and specificity towards arabitol, cofactor imbalance might have hindered efficient arabinose assimilation. Cofactor engineering will be critical to optimize arabinose assimilation in *Y. lipolytica*.

## CONCLUSION

We identified the two most promising pentose-specific transporters TRP6_Yli_ and TRP22_Yli_ that can enhance xylose consumption in *Y. lipolytica,* by screening a set of 16 putative transporters. Guided by bioinformatic, enzymatic, and transcriptomic analyses, we discovered that *Y. lipolytica* has the dormant native pentose metabolism to assimilate both xylose and arabinose. By overexpressing the native XYL2_Yli_ and TRP6_Yli_ (or TRP22_Yli_), it is possible to active this dormant pentose metabolism. Shared transporters and intermediate enzymes such as XYL2_Yli_ (also ADH_Yli_) can negatively affect efficient assimilation of pentose sugars. Due to low activity of arabitol dehydrogenase (ADH_Yli_), it is considered to be the rate-limiting step and makes arabinose assimilation xylose-dependent. Overall, this study sheds light on the cryptic pentose metabolism of *Y. lipolytica* and further helps guide strain engineering of *Y. lipolytica* for enhanced assimilation of pentose sugars.

## ACKNOWLEDGEMENT

This research was funded by the National Science Foundation (NSF CBET#1511881).

